# PaintOmics 3: a web resource for the pathway analysis and visualization of multi-omics data

**DOI:** 10.1101/281295

**Authors:** Rafael Hernández-de-Diego, Sonia Tarazona, Carlos Martínez-Mira, Leandro Balzano-Nogueira, Pedro Furió-Tarí, Georgios J Pappas, Ana Conesa

## Abstract

The increasing availability of multi-omic platforms poses new challenges to data analysis. Joint visualization of multi-omics data is instrumental to understand interconnections across molecular layers and to fully leverage the biology discovery power offered by the multi-omics approach.

We present here PaintOmics 3, a web-based resource for the integrated visualization of multiple omic data types onto KEGG pathway diagrams. PaintOmics 3 combines server-end capabilities for data analysis with the potential of modern web resources for data visualization, providing researchers with a powerful framework for interactive exploration of their multi-omics information.

Unlike other visualization tools, PaintOmics 3 covers a complete pathway analysis workflow, including automatic feature name/identifier conversion, multi-layered feature matching, pathway enrichment, network analysis, interactive heatmaps, trend charts, etc. It accepts a wide variety of omic types, including transcriptomics, proteomics and metabolomics, as well as region-based approaches such as ATAC-seq or ChIP-seq data. The tool is freely available at http://bioinfo.cipf.es/paintomics/.

## INTRODUCTION

The increasing popularity of multi-omic experiments motivates the need for developing tools to combine the multi-layered measurements in the same integrative analysis. The heterogeneity, high dimensionality and multiple interconnectivity of the multi-omics data are distinct aspects that require special attention in this type of studies. While different analysis strategies exist for multi-omics feature selection and for the discovery of relationships among them (1, 2), graphical visualization of the combined datasets is a general and valuable way of simplifying information and assisting interpretability (3).

Several resources for integrative visualization are already available in the context of systems biology. Two well-known tools for graph analysis and visualization are Cytoscape (4) and Gephi (5). These applications provide numerous functions for exploring, manipulating and analyzing complex networks and are complemented with many plugins that enable specialized analysis of molecular data (3). Similarly, the web-based workbench VisANT (6) includes several tools for drawing and analyzing large biological networks and the ability to combine multiple types of networks to systematically analyze correlations with the phenotype. Another interesting tool is 3Omics (7), a web application specifically designed for the analysis of human data, which supports transcriptomics, metabolomics and proteomics datasets. Using 3Omics, users can perform correlation analysis, co-expression profiling, phenotype mapping, pathway and GO enrichment analysis on each dataset, and visualize results graphically.

Alternatively, other software solutions explore interactions among biological features assisted by existing knowledge, usually curated, such as metabolic or signaling pathways. Pathways are a fundamental part of interpreting omics data, as they provide the biological context for a given observation (8). One popular tool for pathway-based visualization is MapMan (9), that allows large datasets, including multiple conditions or time-series experiments, to be displayed as pathway diagrams. Another example is KaPPa-View (10) a web-based tool for integrating transcript and metabolite data into pathway maps. Luo and Brouwer introduced Pathview (11), an R/Bioconductor package for data integration and visualization using KEGG pathways, which has recently been launched as a web tool (12). Pathview allows integration of a wide variety of biological data based on pathways analysis, as long as the omics features are previously mapped to genes. Finally, Garcia-Alcalde et al. developed PaintOmics 2 as a web-based tool for integrated visualization of transcriptomics and metabolomics data using KEGG pathways as a template (13).

Despite being useful, these tools do have some limitations in terms of effective data integration and visualization. As a general rule, network-based tools are useful for identifying the interconnections between multiple biological features, but the size and complexity of the network allied with loose integration to existing knowledge (e.g. pathway diagrams) often hamper interpretation. Conversely, pathway-based solutions reduce the size of the displayed data by grouping the information based on previous knowledge, but they do not allow new molecular connections to easily be inferred. Moreover, most of the tools do not allow researchers to easily integrate data from chromatin profiling experiments or regulatory elements such as microRNAs (miRNAs) or transcription factors (TFs).

Here we introduce PaintOmics 3, a web-based application for integrative visualization of multiple biological datasets on KEGG pathway diagrams. As opposed to other visualization tools, the system covers a complete multi-omics pathway analysis workflow, including support for data from virtually any type of omics measurement, automatic feature name/identifier conversion, pathway enrichment and network analysis. PaintOmics 3 implements the latest technologies in web-based visualization, providing powerful exploratory tools of complex data, conveying to explanatory views of biological processes.

## MATERIALS AND METHODS

### The PaintOmics 3 architecture

The PaintOmics 3 platform is a Client-Server web-application developed entirely using open-source resources (Figure 1). The server side is built using Python 2.7 (14) and R (15) languages and is in charge of processing remote user requests, managing access to biological information stored in a non-relational database implemented in MongoDB (16) and performing a variety of statistical and bioinformatics analyses. The client side is responsible for the data presentation to the user web-browser and was entirely developed using JavaScript language and HTML5 technologies. Communication between the client applications and the server-end is handled by AJAX mechanisms where data are exchanged encoded using JSON.

**Figure 1.**
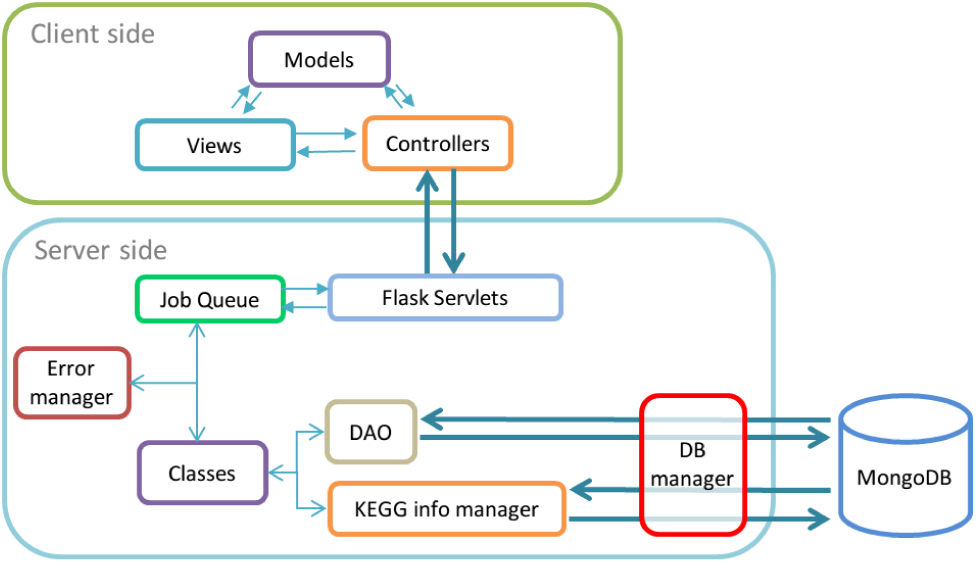
Overview of the PaintOmics 3 architecture. The platform follows the Client-Server paradigm. The Client side implements the Model-View-Controller pattern. The Server side is divided in several subcomponents. The main entry point for the client’s requests are Flask servlets, that manage specific tasks. Heavy computational requests are encapsulated in *job* objects and queued for execution as soon as enough resources are available. Interaction with the database is made through Data Access Objects (DAO) and connections are controlled by a database manager.

### Input data

Input data for PaintOmics 3 are processed feature-level measurements in a tab-delimited file for each omic type. When available, a second file can be provided with a list of relevant features, for instance a list of differentially expressed (DE) genes (Supplementary Figure 1). Experimental conditions should be the same across omics. Ideally, the quantification file should contain one column per experimental condition, so replicated measurements should be averaged. To maximally benefit from the coloring rules implemented in the tool, quantification values should be provided as the log fold change between a case condition and a control/reference condition. The omic data types accepted by PaintOmics 3 can be broadly classified into four categories:

#### Gene-based omics

The biological features measured are, or can be, translated into genes. Some examples are mRNA-seq or microarrays, where measurements are made at the gene or transcript level, and proteomics, where protein quantification can be imputed to the respective coding gene. PaintOmics 3 accepts Entrez Gene IDs as gene identifiers, but it also includes a module for Name/ID translation that allows users to input many other identifiers or naming domains. This module fetches the translation information from public databases such as Ensembl, PDB, NCBI RefSeq, and KEGG, whenever they are available, greatly facilitating data input.

#### Metabolite-based data

The biological features are metabolites. PaintOmics 3 includes some tools for resolving any existent ambiguity when matching user’s metabolites to KEGG compound names. For each input metabolite, it generates a list of potentially related compounds that are ranked with a score based on the similarity of names, and users can manually review, and eventually change, metabolite assignments (Supplementary Figure 2).

**Figure 2.**
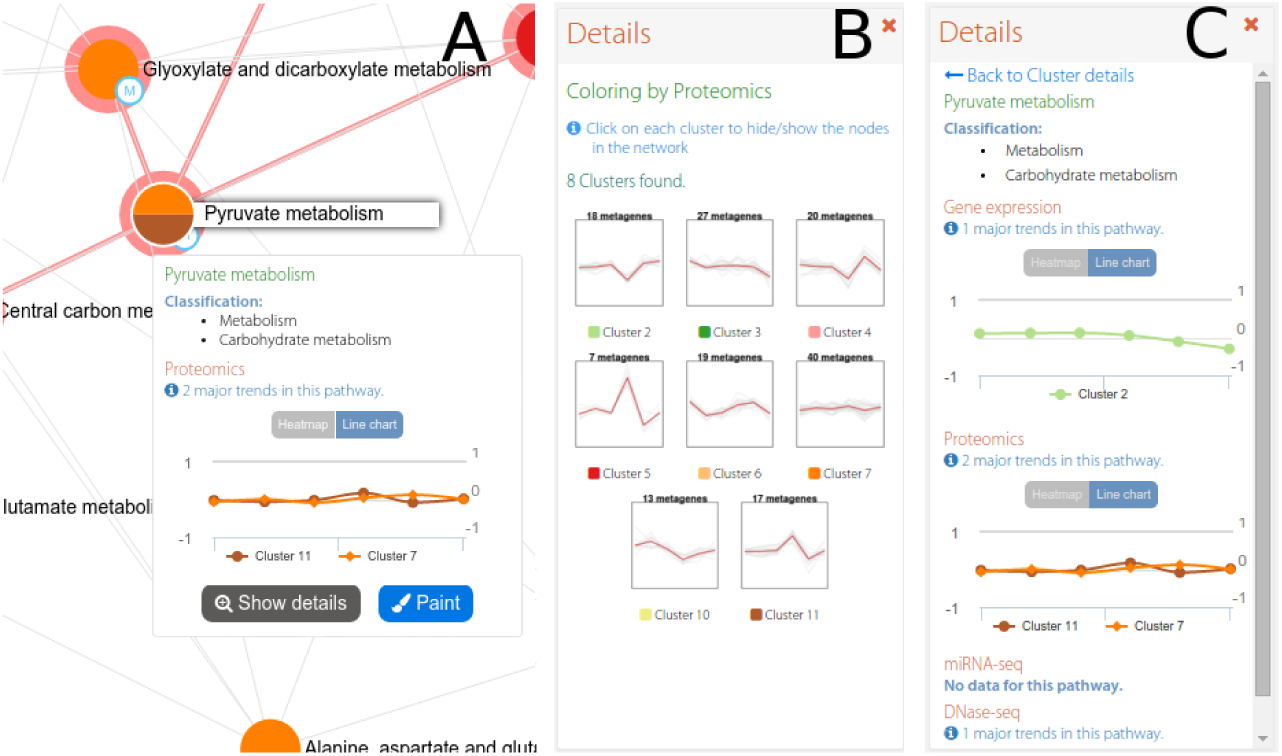
The interactive pathway network in PaintOmics 3. The interactive network panel **(A)** is complemented by a secondary panel showing the trends for all pathway clusters in a given omic **(B)**, or the trends for each omic in the chosen pathway **(C)**.

#### Region-based omics

The measured features are a set of genomic regions, such as those obtained by ChIP-seq, DNase-seq, ATAC-seq, Methyl-seq, etc. For omics in this category, chromosome, start and end position, and a quantification value for the regions must be provided, together with a GTF file with the reference genome annotation. PaintOmics 3 maps each region to its proximal gene(s) with the RGmatch tool (17), which takes into account the region relative position with respect to the specific areas of the gene (promoter region, first exon, intronic areas, etc.). Also, the users can provide regions of interest for their study.

#### Regulatory omics

This category basically refers to miRNAs or transcription factors. Since they are regulators of gene expression, they are mapped to genes. For that, PaintOmics 3 requires an additional tabulated file containing a mapping between miRNA/TF → target gene associations. The system processes the input quantification data and assigns the expression values to the known list of target genes for each regulator.

PaintOmics 3 can process multi-omics datasets with any combination of omics platforms.

### Pathway analysis options

Once data are submitted PaintOmics 3 maps all features to the KEGG database and returns a pathway analysis interface where users can easily keep or hide pathways for downstream analysis, explore pathway networks at different omic layers, visualize single and joint pathway enrichment results, and access specific pathways for interactive browsing.

#### Pathway enrichment

For each omic dataset, and provided that a list of relevant features is included, PaintOmics 3 performs a single Pathway Enrichment Analysis (PEA) using the Fisher’s Exact test. Both p-value and false discovery rate (FDR) adjusted p-values are returned for each dataset. A joint pathway enrichment p-value considering all available omics data is also computed by applying either the Fisher combined probability test (18) or the Stouffer’s method (19). The latter introduces a weight for each individual p-value that allows to control the contribution of each omic type to the computation of the joint enrichment. This is interesting as not all platforms have the ability to measure molecular features with the same comprehensiveness. By default, PaintOmics 3 assigns a weight to each omic that is the percentage of features mapped to pathways in that omic.

#### The multi-omic pathway interaction network

PEA results presented as a list of pathways often fall short to reveal relationships between cellular processes and to offer comprehensive representations of biological systems. PaintOmics 3 includes a tool to create *pathway networks* based on the multi-omics data. In this network, nodes represent pathways and edges indicate shared features among them or KEGG database connections. Each pathway in the network is summarized by one (in some cases several) representative profile obtained by dimension reduction techniques, that recapitulate the major behavior of the pathway along the conditions of the study (20). These pathway profiles are then used to obtain clusters of pathways with similar trends, and the pathway network is colored according to this clustering (Figure 2). In this way, pathways with the same pattern of change can be easily grouped and, if also connected by edges, their molecular relationships revealed. Pathway profiles can be obtained for any of the available omics data and hence networks can be built from each molecular layer perspective. The pathway network tool also includes several options for node selection based either on static (KEGG database) or dynamic (experimental) data.

#### Multi-omic visualization of single pathways

One of the core features of PaintOmics 3 is the visualization of user input data onto individual KEGG pathways. Figure 3 illustrates an example of the typical workspace for pathway exploration. The main panel contains the pathway diagram colored according to the input data. Users can easily navigate through this panel and visualize the different values associated with each biological feature mapped on the KEGG map (Figure 3-A), Feature (genes or metabolites) boxes are divided in as many sections as columns in the input files, and in up to three rows to display different omics measurements. Boxes are colored based on a blue (low) to red (high) gradient scale calculated on each omics range of values after discarding extreme observations. Relevant features are highlighted by a thicker border and a special mark at the top right corner. Two additional panels are available at the pathway workspace. The *Pathway Information* panel includes summary information for all omics data in the pathway, including the major trends for each type of measurement (Figure 3-C). The *Global Heatmap* panel displays the pathway features values in the form of consecutive heatmaps, one per omic dataset (Figure 3-B). More details about the functionalities of this module can be found in Supplementary Data.

**Figure 3.**
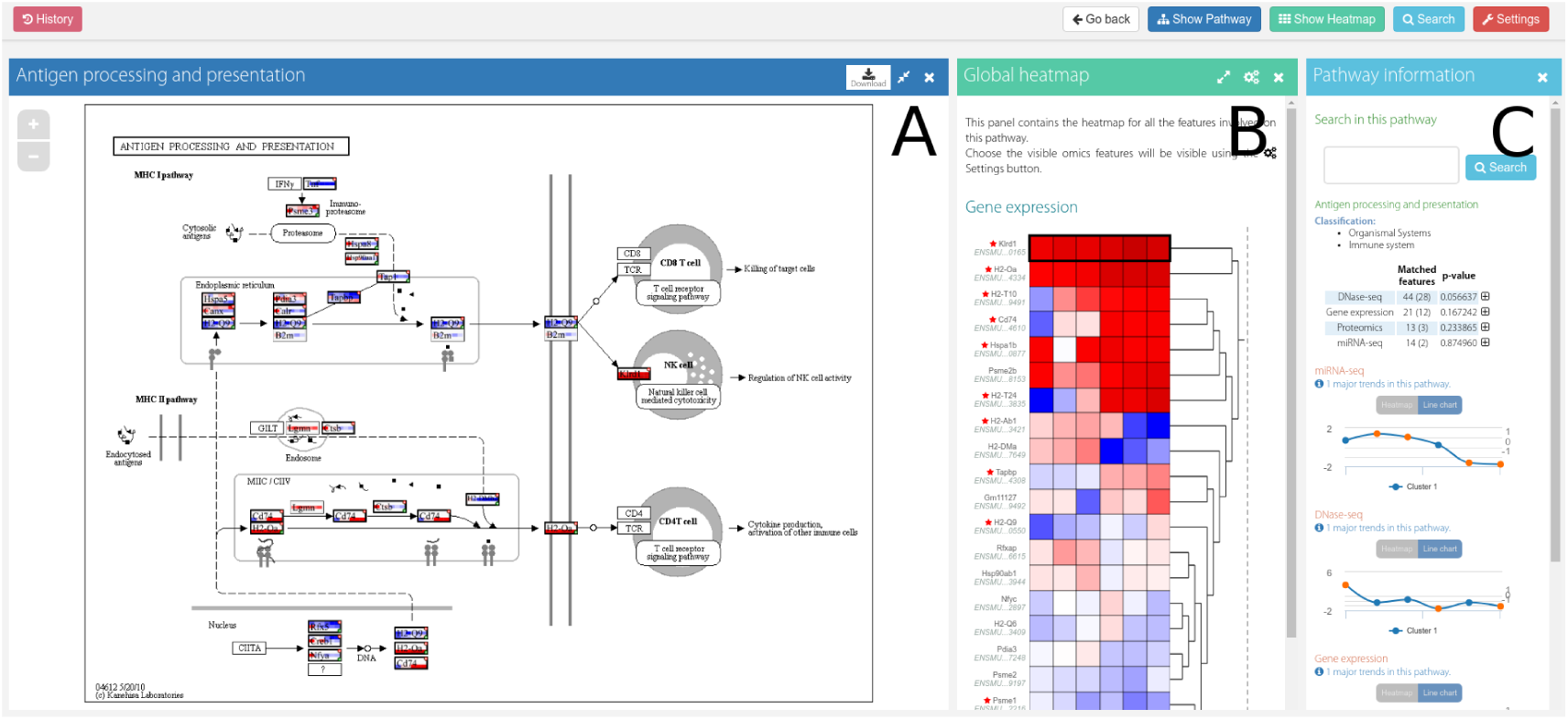
Workspace for pathway exploration in PaintOmics 3. The layout for pathway exploration is divided into three panels. The main panel **(A)** contains the interactive pathway diagram, the *Global Heatmap* panel **(B)** displays multi-omics data in the form of heatmaps, and the *Pathway Information* panel **(C)** contains search and summarizing functions.

### Availability and requirements

PaintOmics 3 is free to use and is distributed under the GNU General Public License Version 3. A public copy of the application is hosted at the CIPF facilities (http://bioinfo.cipf.es/paintomics) and sources are available at GitHub (https://github.com/fikipollo/paintomics3). The documentation and guides for users and administrators are available at the free web platform Read the Docs, and also stored at the GitHub repository. PaintOmics 3 server-side application has been extensively tested on Ubuntu and Debian Linux servers.

## USE CASE

To illustrate PaintOmics 3 use, we selected a multi-omic dataset from Cacchiarelli *et al.* where they analyze changes in gene expression and chromatin state that occur during human reprogramming of immortalized fibroblasts (21). This study includes transcriptomics (RNA-seq and small RNA-seq), methylation (RRBS-seq) and region-based histone modification (H3K4me3 ChIP-seq) data taken at different time points after reprogramming. Details on data preprocessing and input for PaintOmics 3 can be found in the Supplementary Data file.

In order to analyze the Cacchiarelli’s multi-omics dataset with PaintOmics 3, we started by excluding *diseases* related and some *organismal systems* pathways that were considered irrelevant for the study. This resulted in 199 pathways being mapped by multi-omics data, of which 25 were found to be significant on the combined enrichment test (Supplementary Figure 5). Most enriched pathways were regulated at the gene expression and H3K4me3 layers, with a large degree of overlap, which is in agreement with the role as active promoter mark associated to H3K4me3. Enriched pathways at the Methyl-seq and miRNA-seq layers were much fewer, suggesting that these regulatory levels were not as coordinated and ubiquitous.

Significant pathways include hormone and drug detoxification, but mainly signaling, communication and cell lineage. To understand the dynamics and relationships between these processes we analyzed the multi-omic pathway network (Figure 4). The network is dominated by signalling and lineage pathways that are activated between 10 (cluster 5) and 24 days (cluster 6) after reprogramming, including Ca2+ signalling and Wnt signalling (Figure 4-A), also identified by Cacchiarelli *et al.* as transient early upregulation. These pathways are part of H3K4me3 cluster 1 (Figure 4-B), that represents an increase of active promoter histone marks in time. Previous reports indicated that these epigenetic changes enable direct reprogramming to pluripotency (22). On the contrary cluster 4, containing extracellular matrix component and focal adhesion pathways, shows a down-regulation pattern, that was also observed in the original publication and might reflect the need of loose intercellular structures during the reprogramming process.

**Figure 4.**
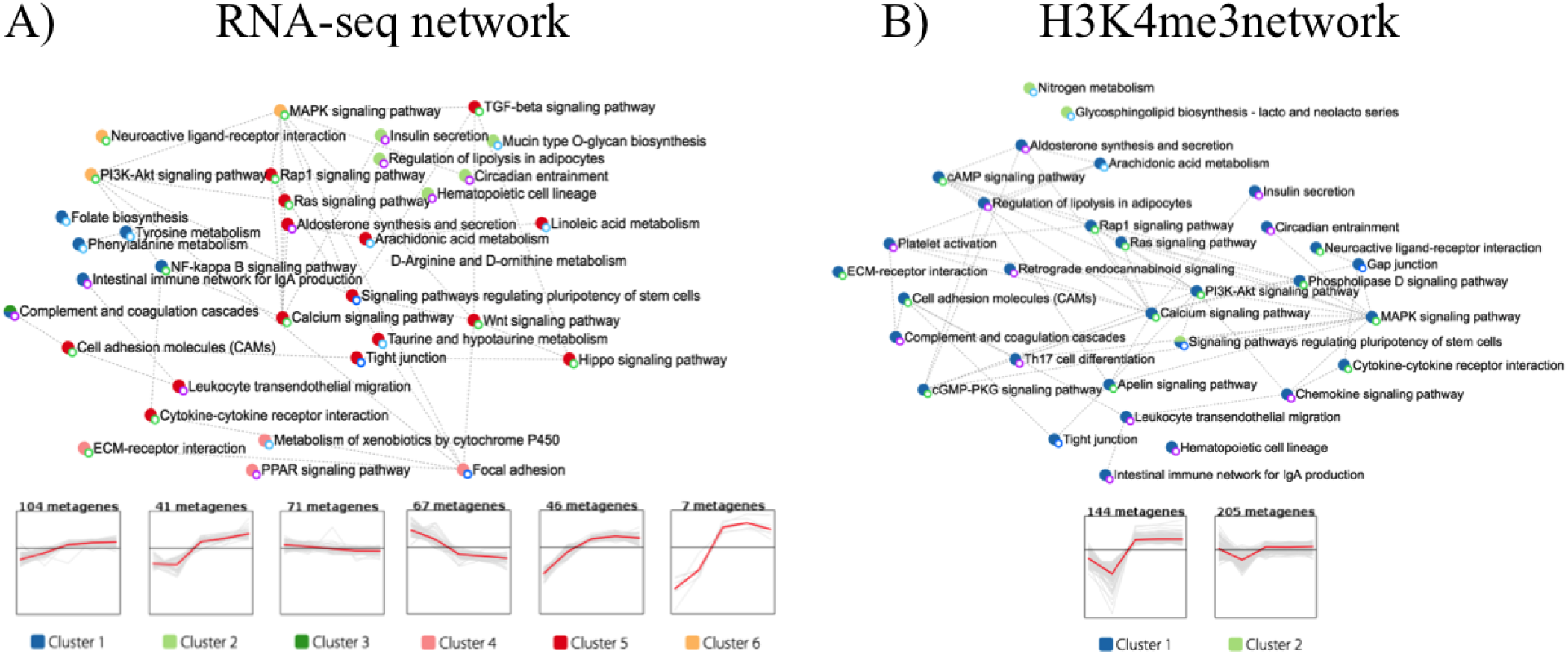
Pathway networks and cluster profiles of representative temporal patterns. Network **A** is colored according to gene expression data. Network **B** is colored according to H3K4me3 ChIP-seq data.

A significant pathway detected in PaintOmics 3 was “Signaling pathways regulating pluripotency of stem cells” (Figure 5). This is a complex pathway in which both naïve and primed stem cells signaling are represented. Major variation trends computed for this pathway by PaintOmics 3 (Figure 5, bottom panels) indicate a general gene expression up-regulation from 10 days after reprogramming that is halted at the last hIPSC stage. H2K4me3 is strong from day 24 while the DNA methylation is consistently down-regulated as reprogramming progresses. The miRNA regulation appears to kick-on at 24 days for genes in this pathway. However, regulatory patterns may vary for different genes. Some examples are shown on the right side of the figure. Most activated genes, including the three key pluripotency genes, POU5F, SOX2 and NANOG are over-expressed and have positive H3K4me3 marks at their promoters (detailed data shown for SOX2). This is in agreement by the activity of several epigenetic regulators like SMARCAD1 and SETDB1, histone modifiers that increase the DNA accessibility at promoters of these genes (23). Other genes in the pathway are down-regulated and PaintOmics 3 reveals potential mechanisms. For example, expression of AKT3 and SMAD3 remain low together with absence of H3K4me3 marks while PIK3R1 is down-regulated with a strong DNA methylation pattern. Interestingly, down-regulation of PIK3R1 has been shown to promote a stem-like phenotype in renal cancer cells through the AKT/GSK3/CTNNB1 pathway, due to AKT phosphorilation (24). Although the AKT genes appear down-regulated in Cacchiarelli’s data, their active involvement in hIPSC has been reported (25). We speculate that the observed lower expression of the AKT isoforms could be compensated by their post-translational activation through the PIK3R1 pathway. Finally, other regulatory patterns appear in this pathway, for example the INHBA gene details box reveals down-regulation is correlated with an strong induction of associated miR-509-3.

**Figure 5.**
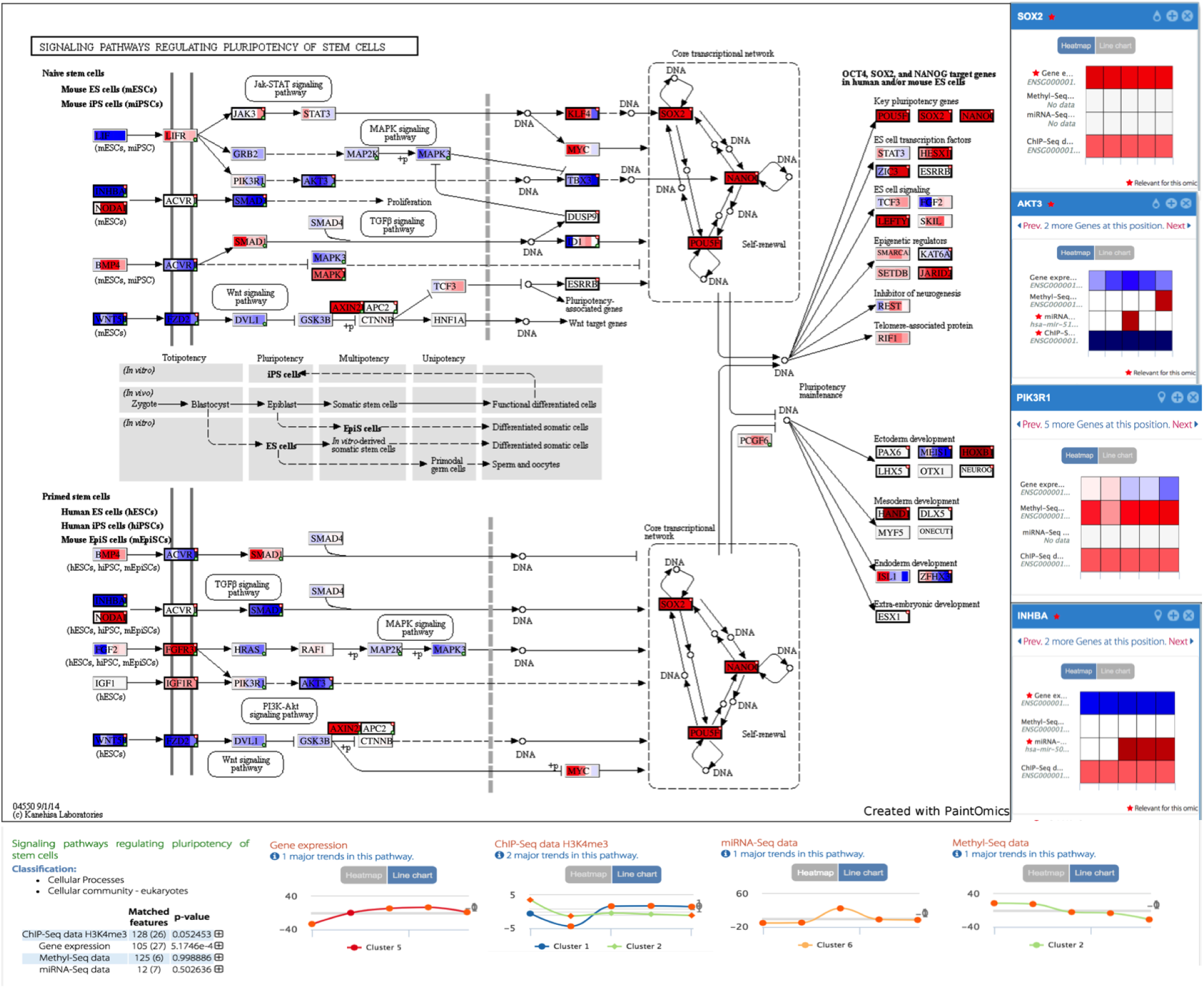
Interactive KEGG diagram for Signaling pathways regulating pluripotency of stem cells. Data obtained from Cacchiarelli’s multi-omics study

## DISCUSSION

We introduce here a novel application for integrative visualization of many different types of omics data, PaintOmics 3, which works as a one-click web tool to enable comprehensive exploration of multi-omics data under the light of pathway models. Using this tool, researchers can easily move through different levels of regulation within biological systems. The combination of a network-based functionality for pathway interactions with they multilayer painted pathway maps leverages the advantages of these two types of visual representations. The network provides an overall view of regulated processes while keeping a manageable size as it uses pathways rather than genes as nodes, and thanks to the filtering options offered by the application. The utilization of omics profiles to represent pathways and coloring the network according to this information is helpful for understanding functional relationships across different molecular layers. The single pathway colored map gives the user a full view of the pathway multi-omics data, while the details panels available for each pathway node facilitates analysis of complex nodes by providing multi-omics values for each of gene independently.

PaintOmics 3 is unique in its strategy to bring data from gene trans- and cis-regulatory factors down to the pathway view. Cis-acting elements will be typically linked to gene nodes thanks to the *Region-based omics* input and PaintOmics 3 will calculate associations directly from bed-like files. This kind of chromatin-based information is normally represented using genome browsers, which completely ignore pathway information and hence cannot study coordinated chromatin changes for cellular processes. Trans-acting regulatory elements (*i.e.* miRNAs, TFs) can be linked through the *Regulatory omics* input. In this case, an association file must be provided by the user. Although this might appear as a limitation, it is in fact an effective solution to provide the tool with full flexibility to visualize any kind of trans-regulatory data the user might have, for example PAR-CLiP or regulatory lncRNA data.

Finally, PaintOmics 3 comes with a wide range of customization and auxiliary functions that allow data and job storage, id conversion, re-coloring, rescaling, filtering, etc. to create in each case the most informative representation of the complex multi-omics dataset. The current version of PaintOmics 3 has been available since 2013 and it has been used by more than 3,000 users world-wide. The variety of organisms supported by PaintOmics 3 outperforms other domain specific tools such as MapMan, KaPPa-View (available for plants) or 3Omics, only available for human, making PaintOmics 3 a highly versatile tool to represent multi-omics information.

